# MutPrior:An Ensemble Method for Ranking Genes in Cancer

**DOI:** 10.1101/058222

**Authors:** Shailesh Patil, Sreya Dey, Randeep Singh

**Affiliations:** SAP labs India Pvt. ltd

## Abstract

Root cause analysis of cancer as well of design of personalized treatment depends on the ability to prioritize mutated genes in cancer. In this paper, we propose a novel approach 'MutPrior' to prioritize genes in a given caner. We hypothesize that a gene is important for cancer if it has high functional impact mutations, is strategically important for network stability and has high relevance to the disease. This approach integrates functional impact scores, centrality in gene-gene interaction network and disease relevance scores to prioritize the mutated genes. MutPrior outputs a prioritization of genes which is more actionable than any current approaches. In the process, we do away with the arbitrary cutoffs as well as confusion caused by notions of driver-passenger.

## I. INTRODUCTION

Carcinogenesis or oncogenesis, the process of formation of cancer cells from normal cells, involves high mutation rates and structural variants of the genomic material. However, only a small subset of these mutations plays an active role in the proliferation and growth of tumor cells. Such mutations are called driver mutations. These mutations can be the root causes and hence, potential drug targets of different cancer types. So, we want to distinguish them from mutations that are not functionally associated with oncogenesis, or passenger mutations. The existing methods for driver mutation detection can be broadly classified into the following three classes: Methods based on Hypothesis Testing, Methods using Machine Learning techniques and Methods based on pathways or gene sets.

Hypothesis Testing based methods find driver genes in a tumor sample by calculating the probability of observing an extreme feature value purely by chance.

Some tools use machine learning mechanisms such as clustering or classification to distinguish driver genes from passenger genes.

Since genes often interact in pathways or as functionally related gene sets, some methods utilize this interaction of gene sets, instead of individual genes, for analyzing driver mutations.

## II. Related Work

### *A.* *Methods using Hypothesis Testing*

In MutSig, genes which are mutated at a rate significantly higher than the Background Mutation Rate (BMR) are detected as driver genes. However, the estimation of BMR is not accurate due to heterogeneity in the mutational processes in cancer. MutSigCV [1] is an improvement on MutSig that accounts for this heterogeneity. Here, genes are mapped to a covariate space and the inter gene distances are used to find the nearest neighbors of each gene. The BMR is then obtained by pooling the nearest neighbors. MutSigCV is adaptive and can be personalized but it does not detect genes with lowly recurrent mutations and has false positives as well.

OncodriveFM [2] identifies genes with accumulation of variations with high functional impact (FI) scores as driver genes. It takes the gene-wise FI scores from SIFT [3], PolyPhen2 [4] and Mutation Assessor (MA) [5], calculates p-values for each tool and combines them using Fisher's combined probability test. Unlike recurrence based approaches, this method is not limited by number of samples and can detect some lowly recurrent genes. However, averaging the FI scores across a gene leads to possible loss of useful information.

OncodriveCLUST [6] works on the hypothesis that gain-of-function mutations always tend to cluster in specific protein position. In this method, positions with mutation rate higher than a Background Mutation Rate are combined and extended to form clusters, which are then assigned scores. It identifies genes mutated in specific protein positions but fails to recognize lowly recurrent mutations and some loss-of-function mutations.

### *B.* *Machine learning techniques*

OncodriveROLE [7] uses a Random Forest classifier to classify driver genes into oncogenes (or Gain-of-Function genes) and tumor suppressor genes (or loss-of-function genes) based on features such as relative abundance of truncating mutations, protein affecting mutations(PAMs) and degree of clustering of missense mutations. Combination of multiple signals controls the rate of false positives identified. However, it fails to identify some lowly recurrent mutations.

CHASM [8] selects features based on Mutual Information and uses Random Forest classifiers to classify mutations into driver and passengers. The fraction of trees voting in favor of the passenger class for a gene is converted to a score, which is in turn assigned a p-value by comparing it to a null distribution formed from synthetically generated passenger mutation dataset. It is one of the first methods to capture large-size custom datasets. However, recent studies have shown that that the occurrence of passenger mutations is affected by factors such as replication time and gene expression. These are not sufficiently captured by a set of random synthetic passenger mutations as used in CHASM.

CanDrA [9] overcomes this drawback by using real passenger mutation data. It uses a two-step feature selection method followed by SVM classification to group mutations into 3 categories: drivers, passengers and no-call. It is cancer-specific. However, it uses its own set of empirical rules to decide driver and passenger mutations while curating the training dataset, which might not be accurate.

### *C.* *Methods using Pathways/Gene Sets*

DriverNet [10] predicts driver genes based on the hypothesis that driver genes are ones with those mutations which will push the gene expression values of the connected genes to some extreme. It uses a greedy algorithm to rank genes and statistical hypothesis testing to compare the results on real data against a randomized dataset. Results revealed a number of rare mutations in known cancer genes typically associated with other cancers. However it has a couple of shortcomings. It does not gracefully handle the directionality of the expression change. Also, the threshold used for determining significant copy number alterations is obtained from third party algorithms and can affect results.

MEMo [11] extracts mutually exclusive gene sets by building event matrix of significantly altered genes and extracting cliques from gene-gene network pairs. This method uses pathway information, resulting in improved accuracy. However it fails to detect co-occurring gene sets. Dendrix [13] identifies mutated driver pathways directly from somatic mutation data collected from large numbers of patients, using a greedy algorithm. It identifies pathways without using any pathway information. However, the greedy algorithm runs the risk of losing out gene sets/pathways that shared some genes with the pathway identified and removed in its previous iteration.

Multi-Dendrix [14] overcomes this by using an Integer Linear Program (ILP) instead of a greedy approach to find multiple driver pathways simultaneously. The results obtained are statistically tested using PPI networks, which themselves contain a large number of false positives.

Tug of War [12] Tug of War models the impact of driver and passenger mutation populations in cancer cells. New traits required for cancer progression are acquired by driver mutations in a few key genes. Most mutations, however, are unimportant for progression and can be damaging to cancer cells, termed passengers. This paper hypothesizes that driver mutations engage in a tug-of-war with damaging passengers. Tug of war model predicts cancer progression.

## III. Need for a new approach

In spite of a plethora of tools available to detect driver genes, the cancer biologists suffer because of the following:

1. The notion of passenger genes is neither well defined nor scientifically established.
2. There is high discordance among the output of multiple tools.
3. Most the algorithms depend of use of background mutation rate, which cannot be accurately estimated.
4. Hypothesis testing based methods assume independence of parameters and fail to exploit correlations
5. Techniques that use machine learning approaches tend to suffer from lack of appropriate labeling of driver vs passenger genes. These tools tend to use a high number of features but often do not have enough training data to deliver desired accuracy.
6. Pathway based approaches rely on accuracy and completeness of interaction network. However, interaction data is incomplete and noisy.
7. All of the current approaches fail to exploit the gene relevance information that is available in literature.
8. Driver detection process employs hard cutoffs and hence might ignore role played by gene below that threshold.

We therefore propose to do away with notion of driver genes. In the next section, we describe a new integrated approach ‘MutPrior that exploits the correlation among the parameters, integrates network and relevance information.

## IV. MutPrior

*We hypothesize that a gene is important for cancer if it has high functional impact mutations, is strategically important for network stability and has high relevance to the disease.*

This hypothesis necessitates an integrated approach to detect genes important in a given cancer. We use an ensemble approach MutPrior for driver detection. We compute scores for genes using several methods and combine them to rank genes by their importance for a given cancer. In a given experimental setup, for each gene, we calculate the functional impact of its mutations and its graph centrality. We also use publicly available relevance score of the genes for cancer. These three scores are combined to obtain final ranking of genes. This final ranking indicates the prioritization of the gene in a cancer. Therefore shortlisting the genes based on cut-offs in not necessary.

### *A.* *Materials and Methods*

In this section, we describe the methods generating various scores and an approach to combine them to generate final ranking. Following are the details of data used in this exercise.

1. We use Chronic Lymphocytic Leukemia(CLL) data from OncodriveFM [2] for generation of functional impact scores.
2. The centrality measure is calculated based on a graph generated from a manually curated Human Reference Network (HRN1) published by MEMo [11].
3. We get disease relevance score from DISEASES [21] method developed by Pletscher-Frankild et. al.

Below, we describe the score generation process ins detail:

### *B.* *Functional Impact Score-Based Method*

This method is based on the hypothesis that genes important for a cancer contain mutations with high functional impact scores. In other words, the important genes are outliers in terms of functional impact scores. As discussed earlier, OncodriveFM is also based on a similar hypothesis. However, it calculates the p-value for each FI score independently and then gets a combined score using Fisher's combined probability test. However, this method violates the independence assumption of the p-values. The FI scores calculated by the tools are not independent. In addition, we believe that there should not be a cutoff on the p-values to shortlist the genes. The score should just be indicators of relative priority of the genes in a given cancer analysis.

Here, we give a brief description of the steps involved in our calculation. This is followed by detailed explanation.

1. Calculate k functional impact (FI) scores for each gene using k known tools of choice.
2. Each gene is represented by a k dimensional vector.
3. Calculate centroid of this data as mean of all the gene vectors.
4. Calculate covariance matrix of this data.
5. Calculate Mahalanobis Distance [19] of each gene from the centroid using above covariance matrix.
6. Sort genes in ascending order of their Mahalanobis Distance.

#### Details

- We calculate functional impact (FI) score of a gene by averaging the FI scores of all the mutations throughout the gene in a sample. The tools used for calculating functional impact scores are SIFT, Polyphen-2 and MutationAssessor. Hence, for each gene we get three functional impact scores.
- We use Mahalanobis distance to detect outliers in the data. This method takes into account the covariance of the gene-wise FI scores calculated by various tools. Let, *g_i_* be the vector of functional impact score for *i^th^* gene, *μ* be the mean FI score vector, and Σ be the covariance matrix of the above data. Then, Mahalanobis distance *d_i_* for gene *g_i_* is calculated as following:

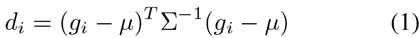
- Most of the genes have very low functional impact scores and hence would be very close to the centroid. As a result, most of the genes will have low Mahalanobis distance scores. Therefore, a gene with very high Mahalanobis distance score would have mutations that have very high functional impact. These genes would be of interest for further study and are highly likely to be driver candidates.

### *C.* *PageRank Score-Based Method*

We try to incorporate the gene-gene interaction information in prioritizing the genes. If genes strategically located in the interaction network are disturbed, they tend to affect larger part of the graph and hence have potential to be relevant for a disease. In terms of graph theory, this translates to genes with high centrality scores. We use PageRank [20] as measure of centrality. Page rank is chosen over closeness and between centrality as it considers not only shortest path and degrees but also captures all the random paths passing through a node. We score each gene as follows:

1. We take the gene-gene interaction network where each gene corresponds to a vertex. Nodes are connected if they are found to interact in the HRN1.
2. We calculate the PageRank [20] score of each vertex (gene), which gives a measure of the centrality of the gene in the network.
3. If genes with high page rank are mutated they might disturb a larger portion of interaction network. Therefore, they might have an important role in the disease.

#### Details

- We use the Human Reference Network used in MEMo as the background gene-gene interaction network graph. It is manually curated and comprises a list of directed interactions between pairs of genes. We have removed self-loops (one gene interacting with itself) from the network and used it for the centrality calculation. Each vertex of the resultant directed, unweighted graph corresponds to a gene and each edge corresponds to an interaction between these two genes in some pathway or biological process.
- Let *G* be a directed graph comprising *n* vertices (*v_1_ v_n_*) and *m* edges (*e_1_ e_m_*). Let A be the transition matrix of *G* where the *A*[*i,j*] denotes the transition probability from vertex *j* to vertex *i*. We define Matrix *M* as follows

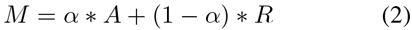

where, *α* is the damping factor, which assumes a value from 0 to 1 (we use *α* equal to 0.85) and *R* is an *n* × *n* matrix with all entries equal to 1/*n*. Let *v* be the *n* × 1 significance vector. Then the sequence, *v, Mv,‥ M^k^ v*, converges to the PageRank score vector.
- Each gene gets a PageRank score that gives a measure of the centrality of the gene in the network. Higher this score, more connected the gene is and disturbance to gene can affect larger part of the network

### *D.* *Text Analysis Based Score*

We incorporate text mining based association score from DISEASES [21] method developed by Pletscher-Frankild et. al. Following is the summary of how these scores were generated by the authors. Abstracts of medical literature are analyzed to find associations or co-occurrences between gene names and disease names. STRING v9.1, which integrates names from Ensembl [23], Uni-ProtKB [24] and HGNC [25], is used for curating human protein and gene names whereas disease names and synonyms are collected from the Disease Ontology [26]. A dictionary is built and a corpus of medical abstracts is analyzed using this dictionary to assign a cooccurrence score to each gene-disease pair. This score takes into account the co-occurrence of the gene-disease pair in the same sentence as well as in the same abstract.

### *E.* *Combining Scores and Ranking Genes*

As discussed, we use 3 scoring schemes to prioritize genes: Scores from FI analysis, Scores from centrality analysis and Scores from text analysis. MutPrior combines the three scores and uses the final score to rank the genes.

1. MutPrior quantile-normalizes each score to make all three distributions similar.
2. MutPrior calculates the geometric mean of the three quantile-normalized scores to get the final score for each gene.

#### Details

- Each score is distributed differently compared to the others. To make the three distributions similar, we use quantile normalization. Quantile normalization works as follows: Let there be three distributions *a*, *b* and *c*. Assume each of them has *n* values. We sort these distributions and calculate the average of the highest values of all three distributions (*avg_1_*), average of the second highest values (*avg_2_*) and so on till the average of the smallest values of each distribution (*avg_n_*). Now in each of the lists, the maximum is replaced by *avg_1_*, second maximum by *avg_2_* and so on. The 3 score lists are normalized as discussed above.
- The geometric mean of the three quantile normalized scores for each gene is the final score of the gene. Genes are sorted by this score and the top scoring genes are selected.

## V. RESULTS AND INTERPRETATION

We verify our results using standard databases like DriverDB [14], Cosmic [15] and Census [16]. We compare top 100 genes in our analysis against the standard databases and 49 genes were present in at least one of the three databases mentioned above. This indicates the effectiveness of our methodology. The genes are listed in Appendix A. A summary of the validation results against the 3 databases are presented in Figure 1. List of top 100 genes is provided in supplementary data.

**Fig. 1.**
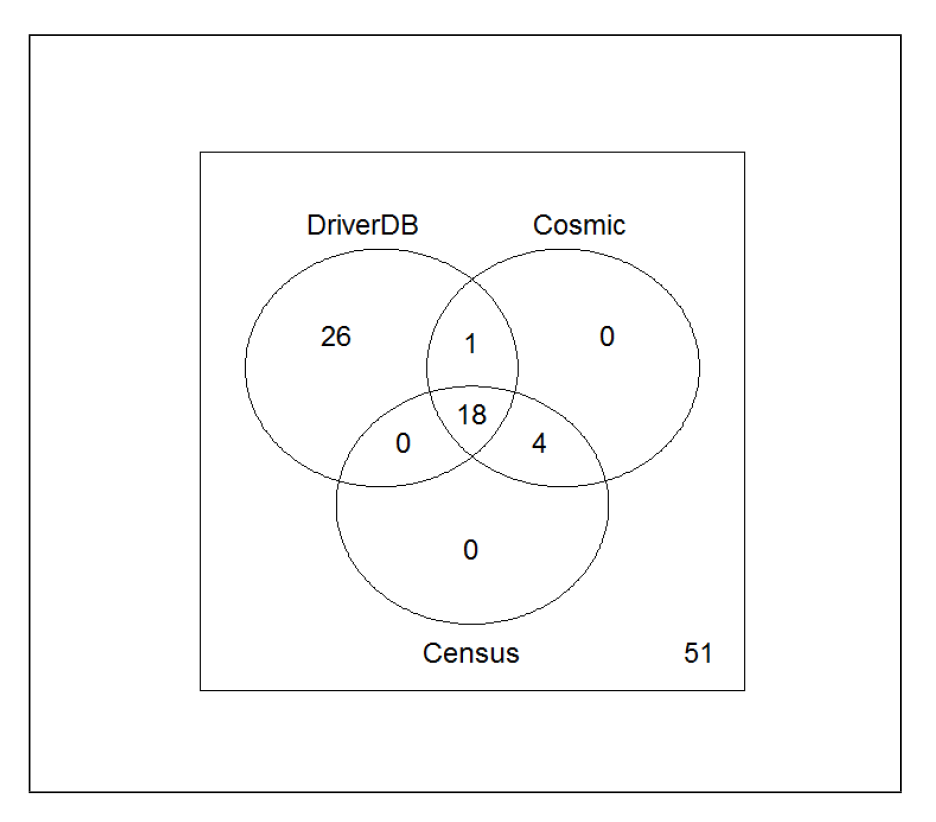
Counts of genes common between top 100 genes from our result and the 3 standard databases

## Acknowledgment

We thank our interns Trenita D,almeida and Abhinav Khare for their assistance with literature survey and collection of data.

